# Short-Term Exposure to Polystyrene Microplastics Alters Cognition, Immune, and Metabolic Markers in an APOE Genotype and Sex-Dependent Manner

**DOI:** 10.1101/2025.05.02.651942

**Authors:** Lauren Gaspar, Sydney Bartman, Hannah Tobias-Wallingford, Giuseppe Coppotelli, Jaime M. Ross

**Affiliations:** Department of Biomedical and Pharmaceutical Sciences, College of Pharmacy, University of Rhode Island, Kingston, RI 02881, USA; George and Anne Ryan Institute for Neuroscience, University of Rhode Island, Kingston, RI 02881, USA

**Keywords:** microplastics, nanoplastics, Alzheimer’s disease, dementia, mouse

## Abstract

Alzheimer’s disease (AD) is one of the most prevalent neurodegenerative disorders and one of the leading causes of death in individuals over the age of 65. Most cases of AD develop sporadically, however, there are several risk factors that have been identified which significantly increases an individual’s risk for developing AD. The most prominent of these is Apolipoprotein E4 (APOE4), which can potentially result in an up to 10-fold greater risk of developing AD. The presence of APOE4 alone, however, cannot be solely responsible for AD as the disease may occur even in the absence of APOE4. Therefore, there must be other contributing factors such as exposure to environmental toxins including heavy metals and pesticides, which have independently been shown to contribute to AD. Nano- and microplastics (NMPs) are plastic particles less than 1 μm and 5 mm in size, respectively, and have only recently been identified as a major environmental pollutant with serious health concerns. Given the adverse health effects that are increasingly being associated with NMPs exposure, we sought to understand how the combination of APOE4 and NMPs exposure may work synergistically to promote cognitive dysfunction and alter key regulatory pathways to impact overall health. Following an acute (3 week) exposure to pristine spherical fluorescently-labeled 0.1 and 2 µm polystyrene (PS) NMPs, we found significant sex-dependent alterations in locomotor and recognition memory in APOE4 mice, but not in APOE3 controls. We additionally found that exposure to PS-NMPs resulted in sex and genotype specific alterations in astrocytic and microglial markers in the brain, and in CYP1A1, a major metabolizer of environmental polycyclic aromatic hydrocarbons, in the liver. These results suggest PS-NMPs may interact with the APOE4 allele to promote cognitive dysfunction and alter immune and metabolic pathways which may contribute to disease-like states.

## Introduction

Alzheimer’s disease (AD) is a multi-faceted neurodegenerative disorder that currently affects nearly 7 million Americans over the age of 65 and is one of the leading causes of death in elderly populations (“Alzheimer’s Disease Facts and Figures,” n.d.)(Zhang et al., 2021)(Liang et al., 2021). Individuals with AD may experience symptoms ranging from memory loss, personality changes, difficulty with problem solving, language impairments, and in severe cases physical symptoms such as loss of bladder control and seizures (“What Are the Signs of Alzheimer’s Disease?,” 2022)(Pinyopornpanish et al., 2022)(Lopez and DeKosky, 2008). These symptoms, which greatly impact the daily lives of those with AD, as well as their caregivers, have prompted AD to become one of the most widely researched age-related disorders. The challenge in researching AD is that most cases are not familial (<5%), but rather sporadic (>95%), meaning there are no clearly inherited genes or mutations that cause AD development (Bali et al., 2012)(Lista et al., 2015). As a result, it is widely believed that there are many contributors to the progression of AD, including genetic risk, socioeconomic, lifestyle, and environmental factors.

Although it has become clear to researchers that no single one of these factors can be attributed with causing AD, there have been several prominent risk factors that have been identified in recent years. One such factor is known as apolipoprotein E4 (APOE4) (Ayyubova, 2024). Apolipoprotein E is a protein that predominantly serves to mediate lipid metabolism and transport (Huang and Mahley, 2014)(Yang et al., 2023). In humans, APOE has been shown to have 3 major isoforms: APOE2, APOE3, and APOE4. APOE3 (Cys112, Arg158) is the most common isoform found in ∼75% of the population and is considered to be neutral towards AD risk. APOE2 (Cys112, Cys158) is the least common isoform found in ∼5% of the population and is considered to be slightly protective against AD development. APOE4 (Arg112, Arg158) is found in ∼20% of the population and is considered one of the greatest known genetic risk factors for AD (Husain et al., 2021)(Wu et al., 2018). Studies suggest that one copy of APOE4 may double an individual’s risk of developing AD, whereas two copies of the allele may increase risk by 10 times or even more (“A rare mutation protects against Alzheimer’s disease, Stanford-led research finds,” n.d.)(Uddin et al., 2019). Additionally, it has been suggested that APOE4 may contribute to earlier onset of AD and increased cognitive impairment as compared to AD patients without APOE4 (M. Di Battista et al., 2016)(Emrani et al., 2020)(Safieh et al., 2019).

While there have been many mechanisms proposed as to how APOE4 contributes to AD including impaired lipid metabolism (Yin, 2023)(Jeong et al., 2019), reduced clearance of amyloid-beta and tau (Prasad and Rao, 2018)(Liu et al., 2017)(Eisenbaum et al., 2024), and induction of neuroinflammation (Parhizkar and Holtzman, 2022)(Dias et al., 2025), there is still no clear mechanism. This is further complicated by the fact that the development of AD can occur in the absence of APOE4. This has prompted researchers to explore how APOE4 may be interacting with other contributing factors of AD to potentially accelerate disease onset and severity. The majority of these studies focus on the combined effects of APOE4 and lifestyle factors, such as high-fat diet (Janssen et al., 2016)(Mattar et al., 2022)(Huebbe et al., 2024) and exercise (Cancela-Carral et al., 2021)(Tokgöz and Claassen, 2021); however, very few have examined the interaction of APOE4 with environmental pollutants, which have independently been shown to contribute to AD pathology. In particular, no study has yet to explore how the presence of APOE4 may increase an individual’s susceptibility to one of the most prevalent emerging environmental pollutants, microplastics (MPs), or how this interaction may accelerate cognitive impairment and other AD pathologies.

Microplastics, defined as any plastic less than 5 mm in size, have recently been identified as one of the most ubiquitous and potentially harmful environmental pollutants (Lim, 2021)(Zhao et al., 2024)(Li et al., 2025). To date MPs have been shown to induce numerous detrimental health effects, including increased oxidative stress (Kadac-Czapska et al., 2024)(Zou et al., 2023)(Jia et al., 2023), increased inflammation (Pulvirenti et al., 2022)(Gaspar et al., 2023)(Luo et al., 2022), alterations in reproductive function (Urli et al., 2023)(Inam, 2025), and gut dysbiosis (Sofield et al., 2024)(Xie et al., 2021). Additionally, MPs have been shown to translocate throughout the body and into tissues such as liver, lungs, heart, and even the brain (Gaspar et al., 2023)(Li et al., 2024). Most recently, it was reported that patients with dementia, on average, have a higher burden of MPs in their brains (Nihart et al., 2025). As such, we sought to explore if the toxic effects induced by MPs could have synergistic outcomes with APOE4 to promote cognitive impairment and other associated markers of neurological decline.

## Materials and Methods

### Animal Husbandry and Microplastics Exposure

Colonies of humanized knock-in APOE3 and APOE4 mice were originally obtained from Taconic Biosciences (Germantown, NY, USA) and the cohorts of female (*n* = 32) and male (*n* = 32) APOE3 and APOE4 mice for this study were bred at the vivarium at the University of Rhode Island. Amongst each genotype and sex, mice were randomly divided between control and exposure groups (n = 8 per group). Starting at 3-6 months of age, control mice received standard drinking water while mice in the exposure group received a 1:1 volume mixture of 0.1 and 2 µm pristine spherical fluorescently-labeled polystyrene nano- and microplastics (PS-NMPs) at a concentration of 0.125 mg/mL (Fig. 1A), as done previously (Gaspar et al., 2023). This concentration, which is higher than the current estimates of human exposure, was chosen to account for the short exposure duration compared to the chronic exposure humans experience in everyday life (Nakat et al., 2023)(Sathish et al., 2020)(Diaz-Basantes et al., 2020)(Senathirajah et al., 2021). Delivery via drinking water was selected over oral gavage to allow for continuous exposure, as well as to minimize external stress that may impact behavior performance. Water bottles were inverted every 10–12 h even though we previously found no particle settlement (Gaspar et al., 2023). Mice were exposed for three weeks, during which time the drinking water was replaced as needed, i.e., every ∼10–12 days. All mice received a standard diet (Teklad Global Soy Protein-Free [Irradiated] type 2920X, Envigo, Indianapolis, IN, USA) and water *ad libitum*. Animals were group-housed by sex with up to 5 mice per ventilated cage with access to a small hut and tissues for nesting. The mice were kept on a 12:12 light: dark cycle at 22 °C ± 1 and 30–70% humidity. Adequate measures were taken to minimize animal pain and discomfort. The investigation was conducted in accordance with the ethical standards and according to the Declaration of Helsinki and national and international guidelines and has been approved by the authors’ institutional review board.

**Figure 1.**
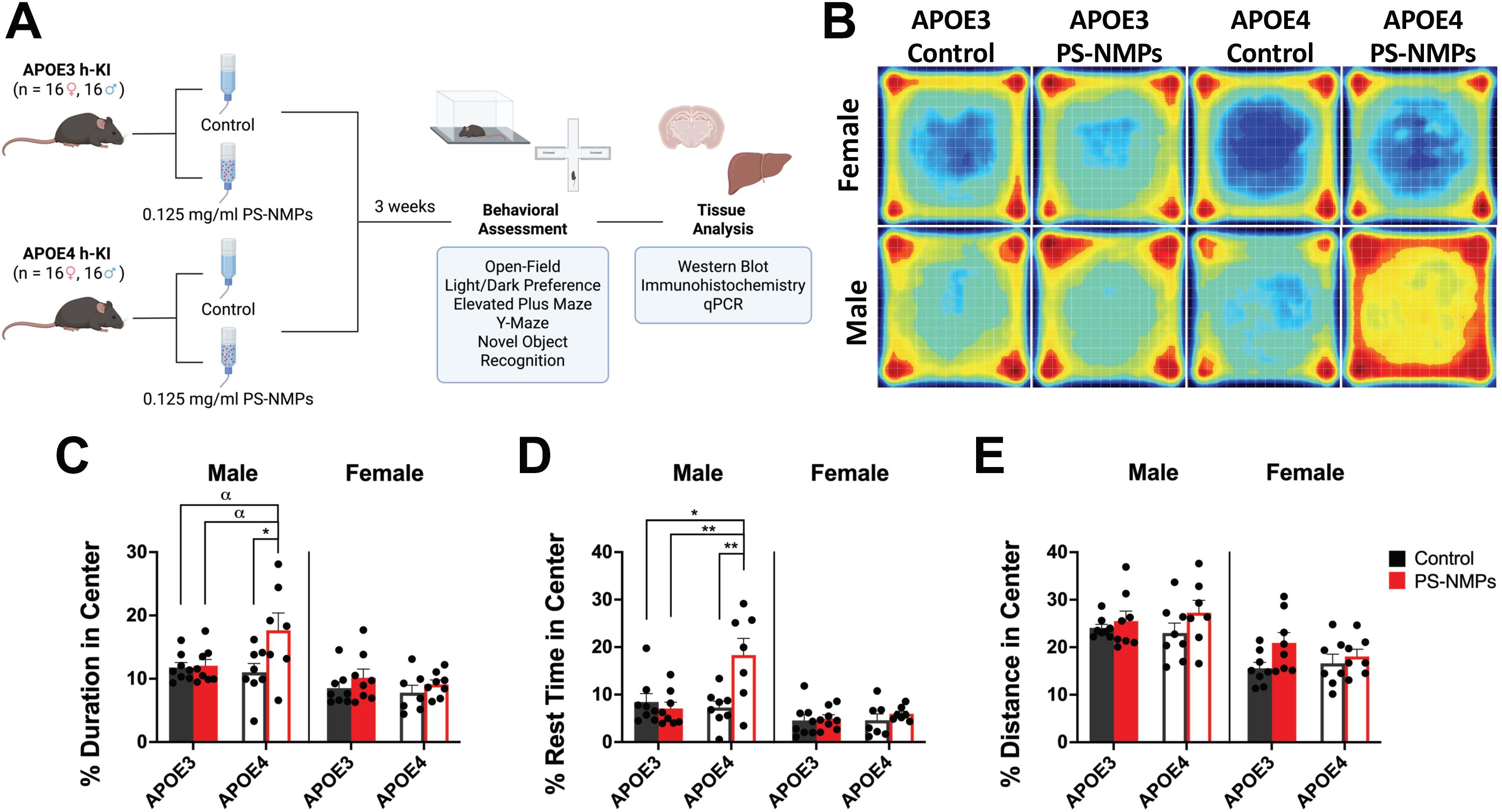
Study design for assessing interplay between APOE genotype and exposure to PS-NMPs and effect of PS-NMPs exposure on spontaneous locomotion in APOE3/4 h-KI mice. **(A)** *In vivo* study design for short-term (3 weeks) exposure to PS-NMPs in 3-6-month-old male and female hAPOE3/4-KI mice (*n* = 8 per group). Schematic was created with BioRender.com (accessed on 29 April 2025). **(B-E)** Spontaneous locomotor activity of 3-6-month-old male and female hAPOE3/4-KI mice (*n* = 8 per group) exposed to PS-NMPs (red), as compared to controls (black). **(B)** Representative heat maps visually demonstrate differences in locomotor activity. Male APOE4 mice exposed to PS-NMPs showed marked increases in **(C)** percent duration in center and **(D)** percent rest time in center. Three-way ANOVA was used to determine the interaction between APOE genotype, sex, and PS-NMPs exposure: **(C**) genotype x sex x exposure p=0.085, **(D)** genotype x sex x exposure p=0.018, **(E)** genotype x sex x exposure p=0.210. Sex-specific significances were determined by two-way ANOVA: **(C_Males_)** genotype p=0.134, PS-NMPs exposure p=0.036, interaction p=0.052, **(C_Females_)** not significant; **(D_Males_)** genotype p=0.022, PS-NMPs exposure p=0.026, interaction p=0.006, **(D_Females_)** not significant; **(E_Males_)** not significant, **(E_Females_)** PS-NMPs exposure p=0.065, (genotype and interaction not significant), with post-hoc analyses, denoted by * *p* < 0.05, ** *p* < 0.01, and α *p* < 0.10 as trending.

### Behavior Experiments

All mice were acclimated in their home cage for 1 h in the testing room and were kept at 22 °C ± 1, 30–70% humidity, and ∼100 lux prior to testing. The testing room was in a neutral, quiet environment, and mice were tested between 9:00 and 17:00 (light phase) by the same researcher, with care taken to stagger the testing of mice from the different cohorts. All apparatuses were cleaned between tests with 70% ethanol. The order of behavior testing are as follows below.

### Open-Field (OF)

An infrared motion detection system (Fusion v6.5 SuperFlex, Omnitech Electronics, Columbus, OH, USA) was used to assess exploratory behavior and spontaneous locomotion. During this assay, the mice were placed in transparent locomotor chambers (40 × 40 × 30 cm) with a grid of infrared beams at floor level and 7.5 cm above floor level for 90 min while their movements were monitored in 5 min intervals in the x-, y-, and z-planes. All horizontal and vertical movements were recorded, and data were analyzed using the Fusion v6.5 software system. All test chambers were cleaned with 70% ethanol after each test period.

### Light-Dark Preference Test (LD)

All mice were placed in locomotor chambers (40 × 40 × 30 cm) divided into light and dark zones that were equipped with a grid of infrared beams at floor level and 7.5 cm above floor level, to assess exploratory and anxiety-related behaviors. All mice were tested for 30 min with their movements being monitored in the x-, y-, and z-planes using an infrared activity-monitoring cage system (Fusion v6.5 SuperFlex, Omnitech Electronics, Columbus, OH, USA). All movements were recorded, and data were analyzed using the Fusion v6.5 software system. Locomotor boxes and inserts were cleaned with 70% ethanol after each test period.

### Elevated Plus Maze (EPM)

The elevated plus maze (EPM) consists of two open (30 × 7 cm) and two enclosed (30 × 7 × 15 cm) arms crossed to form a plus-sign shape and was used to assess anxiety-like behavior. Mice were transported to and from the maze in a darkened container to minimize external stress and stimuli. During the maze, mice were placed on the apparatus (CleverSys, Reston, VA, USA) and allowed to explore for 5 min at ∼300 lux while their movements were monitored via a camera tracking system (Anymaze, Stoelting, Wood Dale, IL, USA). The maze was cleaned with 70% ethanol between each test.

### Y-Maze

To assess short-term memory, the Y-maze, which consists of three identical enclosed arms (30 × 7 × 15 cm) arranged in a Y-shape, was used. Mice were transported to and from the maze in a darkened container to minimize external stress and stimuli. All mice first completed a familiarization trial during which the mice were placed in one arm of the apparatus (CleverSys, Reston, VA, USA) facing the center and were allowed to explore two of the arms for 3 min at ∼300 lux while the third arm remained closed off. The mice were then returned to the home cage for 15 min. After this time, the mice were placed back in the apparatus and allowed to explore again for 3 min with access to the novel arm. All animal movements were monitored via a camera tracking system (Anymaze, Stoelting, Wood Dale, IL, USA). The apparatus was cleaned with 70% ethanol between each test.

### Novel Object Recognition Test (NOR)

To assess spatial and recognition memory, an infrared motion detection system (Fusion v6.5 Superflex, Omnitech Electronics, Columbus, OH, USA) containing a grid of infrared beams at floor level and 7.5 cm above floor level was used to monitor animal movements in x-, y-, and z-directions. During this test, mice were placed into the transparent chambers (40 × 40 × 30 cm) containing two novel objects for 30 min (habituation phase) while their movements were monitored in 1 min intervals. The mice were then returned to the home cage for 3 h. After this time, the mice were placed back in the chamber, where one object has been replaced, for 30 min while their movements were monitored in 1-minute intervals (novel versus familiar object testing). All data were analyzed using the Fusion v6.5 software system. All chambers were cleaned with 70% ethanol after each test period.

### Tissue Preparation

All mice were anesthetized after the behavioral testing was complete with sodium pentobarbital (200 mg/kg) intraperitoneally followed by cervical dislocation. Brain, lungs, heart, liver, kidneys, gastrointestinal tract (GI), spleen, and gastrocnemius muscle tissues were dissected and either post-fixed in 10% formalin (Epredia, Portsmouth, NH, USA) for 24 h at 4 °C, followed by 30% sucrose (*w*/*v*) in 1X PBS, or quickly frozen on dry ice and stored at −80 °C.

### Fluorescent Immunohistochemistry (IHC)

For determining GFAP and IBA1 expression, frozen and embedded post-fixed brain samples from control and PS-NMPs exposed APOE3 and APOE4 males and females were cryosectioned (Leica BioSystems, CM1950, Wetzlar, Germany) at 30 μm taken at −21 °C; free-floating sections were collected into 12-well plate netwells (3477, Corning, Corning, NY, USA) filled with PBS using previously reported methodology (Potts et al., 2020). Briefly, sections were blocked in TBS with 3% horse serum and 0.3% Triton X-100 at RT for 30 min on an orbital shaker and incubated overnight at 4 °C on a rotator with rabbit anti-GFAP primary antibody (1:2000, PA1-10019, Invitrogen, Waltham, MA, USA) and goat anti-IBA1 primary antibody (1:500, 011-27991, FujiFilm Wako, Richmond, VA, USA) in TBS with 1% horse serum and 0.3% Triton X-100. Sections were incubated the following day with donkey anti-rabbit Alexa 488 (1:500, A21202, Invitrogen, Waltham, MA, USA) and donkey anti-goat Alexa 594 (1:500, AB150132, Abcam, Cambridge, UK) secondary antibody in TBS with 1% horse serum and 0.3% Triton X-100 at RT for 2 h on an orbital shaker, while being protected from light. This was followed by hoescht (1:2000) staining. Sections were mounted onto slides (VWR Colorfrost®Plus, Radnor, PA, USA), coverslipped with aqueous mounting medium, and dried at RT for 15–20 min protected from light. Fluorescence imaging (Leica THUNDER DMi8 3D Fluorescence Imaging System, Leica Biosystems, Wetzlar, Germany and LAS X 3D Analysis Software v. 2018.7.3, Leica Biosystems, Wetzlar, Germany) was used to evaluate GFAP and IBA1 expression.

### Western Blot

Fresh frozen brain and liver samples from both sexes and exposure groups for APOE3 and APOE4 mice were lysed in RIPA buffer (50 mM Tris-HCl pH 7.4, 150 mM NaCl, 0.5% deoxycholic acid, 0.1% sodium dodecyl sulfate, 2 mM EDTA, 1% Triton X-100) containing a proteinase (1:100, 78,438, Halt, Thermo Scientific, Fremont, CA, USA) and phosphatase (1:100, P0044, Sigma Aldrich) inhibitor cocktail. Samples were incubated on ice for 30 min and then sonicated (QSonica, Newtown, CT, USA) for 3 min (30–30 pulse) at 4 °C with 30% amplitude and centrifuged at 10,000× *g* for 10 min. Protein concentration was determined using a BCA protein assay kit (23,225, Thermo Scientific, Fremont, CA, USA) according to the manufacturer’s instructions. Lysates were mixed with loading buffer (1610747, Bio-Rad, Hercules, CA, USA) plus 100 mM DTT and incubated at 98 °C for 10 min. Samples were fractionated in an SDS-PAGE precast 8–16% or 4-20% gradient gel (5671104 or 5671094, Bio-Rad, Hercules, CA, USA) and blotted on a 0.45 μm nitrocellulose membrane (10600002, Amersham Protran, Chicago, IL, USA) using transfer buffer (25 mM Tris-HCl pH 8.3, 190 mM glycine 20% methanol). The membrane was blocked (0.1% TBS-Tween with 5% skim milk) and probed with rabbit anti-GFAP primary (1:10,000, PA1-10019, Invitrogen, Waltham, MA, USA) or rabbit anti-CYP1A1 primary (1:500, 13241-1-AP, Proteintech, Rosemont, IL, USA) and goat anti-rabbit-HRP secondary antibodies in 0.1% TBS-Tween (1:3000, 1706515, Bio-Rad, Hercules, CA, USA); then, immunocomplexes were detected by chemiluminescence (Clarity ECL, Bio-Rad, Hercules, CA, USA) and visualized (ChemiDoc, BioRad, Hercules, CA, USA). Images were processed and quantified using appropriate software (FIJI v2.1.0/1.53c, Madison, WI, USA) (Schindelin et al., 2012).

### Quantitative Polymerase Chain Reaction (qPCR)

Samples of liver tissue (30–50 mg) from both sexes and exposure groups for APOE3 and APOE4 mice were lysed, and RNA was extracted and purified (Zymo Direct-zol RNA MiniPrep Plus Kit R2070, Zymo Research, Irvine, CA, USA) according to the manufacturer’s instructions. RNA concentration was determined using a spectrophotometer (NanoDrop, ND-2000, Thermo Scientific, Fremont, CA, USA), and reverse transcription was run (Lunascript® RT Supermix Kit E3010, New England BioLabs Inc., Ipswich, MA, USA) according to the manufacturer’s protocol. Once the cDNA was synthesized, qPCR reactions were run using a Taqman-based master mix (4444557, Applied Biosystems, Waltham, MA, USA), a CYP1A1 primer (Mm00487218_m1, ThermoFisher Scientific, Waltham, MA, USA), and an Actin primer (Mm00607939_s1, ThermoFisher Scientific, Waltham, MA, USA) according to the manufacturer’s instructions using appropriate instrumentation (Roche Lightcycler 96, Basel, Switzerland) and the following conditions: 95°C for 20 s, 95°C for 3 s, 60°C for 30 s x40. The results were analyzed using the appropriate software (GraphPad Prism v. 10, San Diego, CA, USA).

### Statistical Analysis

Data are presented as mean values (M) with SEM. Statistical analyses using unpaired t-tests, one-way ANOVAs, or two-way ANOVAs with Tukey post-hoc multiple comparisons were performed using appropriate software (GraphPad Prism v.10, San Diego, CA, USA). Significances are denoted as \**p* < .05, \*\**p* < .01, \*\*\**p* < .001, and \*\*\*\**p* < .0001, with α *p* < .10 as trending.

## Results

### Alterations in Cognition in APOE4 Mice in Response to PS-NMPs Exposure

After 3 weeks of exposure to PS-NMPs, (Fig. 1A) APOE3 and APOE4 h-KI mice underwent the following behavioral testing, beginning with the least amount of handling: open-field, light-dark preference, elevated plus maze, y-maze, and novel object recognition. To assess spontaneous locomotor activity, we conducted an open-field assay where mice were allowed to explore a dimly lit chamber for 90 minutes while their movements were monitored in x-, y-, and z-directions. We assessed several parameters including duration, rest time, and distance in the center of the chamber (Fig. 1B-E). Male APOE3 mice showed little to no behavioral response to exposure to PS-NMPs in open-field; however, male APOE4 mice exposed to PS-NMPs exhibited striking increases in percent duration in the center (Fig. 1C) and percent rest time in the center (Fig. 1D) following exposure, as also depicted in the movement heatmaps (Fig. 1B). For female mice, no significant changes were observed in these parameters regardless of APOE genotype or PS-NMPs exposure; however female APOE3 mice exposed to PS-NMPs did show a moderate increase (p = 0.1458) in distance traveled following PS-NMPs exposure (Fig. 1E), which recapitulates findings from our previous study in wildtype C57BL/6J females (Gaspar et al., 2023).

To determine if exposure to PS-NMPs affected anxiety-like behaviors in APOE3 and APOE4 h-KI mice, both light-dark preference assay and elevated plus maze (EPM) were conducted (Fig. 2). For the light-dark preference assay, animals were placed in a chamber divided into light and dark zones for 30 minutes with movements monitored in x-, y-, and z-directions, and the duration, distance, and rest time in the light zone were measured. We found that APOE genotype and exposure to PS-NMPs showed no significant effect on any of these parameters in either sex (Fig. 2A-C). During the EPM assay, mice explored two open and two closed arms arranged in a “plus” shape, elevated from the floor, for 5 minutes. Their movements were monitored, including distance and bouts (entries) in open arms. Similar to the light-dark preference test, we found no significant differences in EPM behavior between APOE genotypes or following PS-NMPs exposure in either sex (Fig. 2D, E).

**Figure 2.**
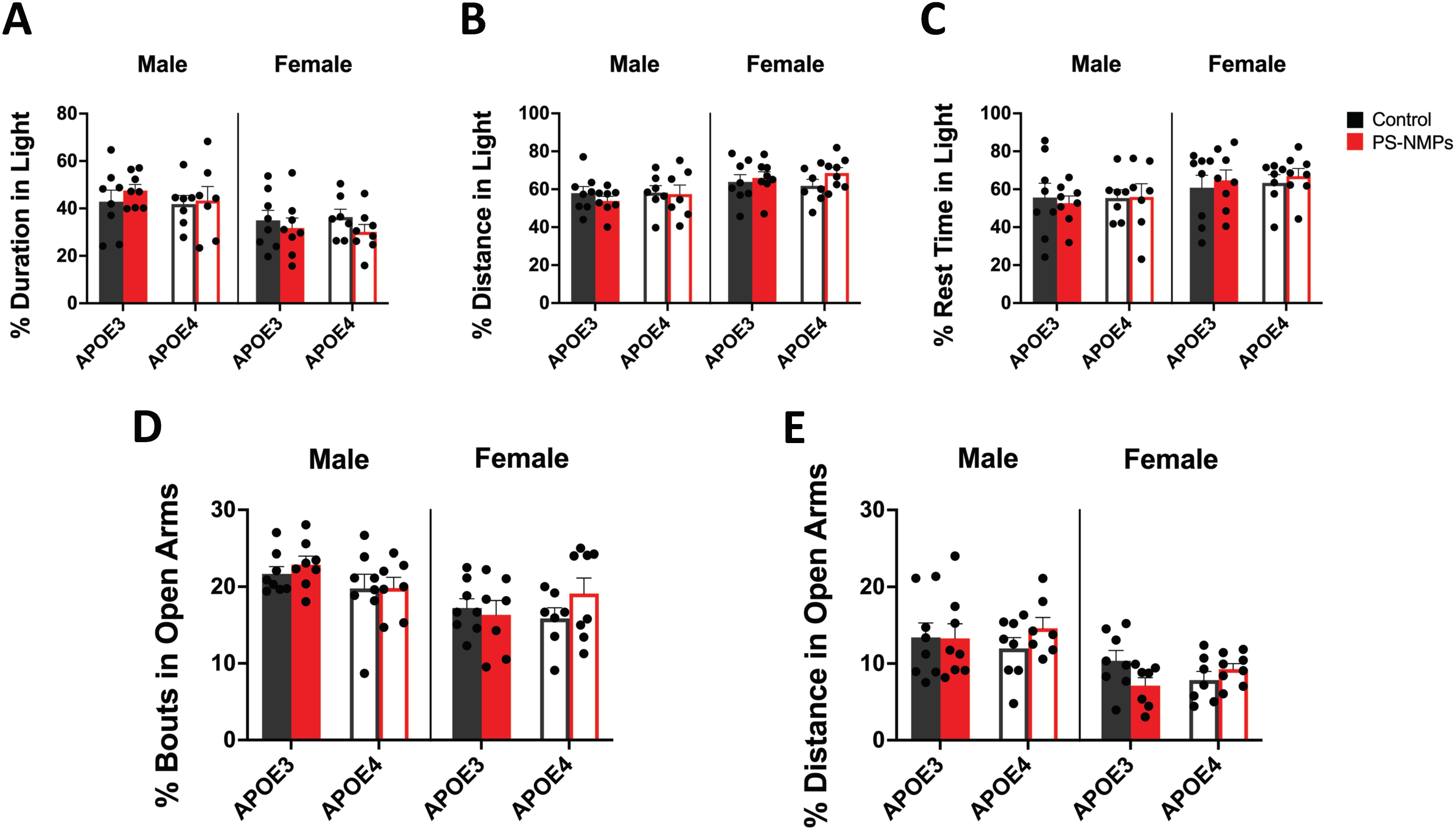
Effect of APOE genotype and exposure to PS-NMPs on anxiety-like behavior. **(A-C)** Light-dark preference of 3-6-month-old male and female hAPOE3/4-KI mice (*n* = 8 per group) exposed to PS-NMPs (red), as compared to controls (black). No significant differences in **(A)** percent duration in light, **(B)** percent distance in light, and **(C)** percent rest time in light were found between genotype or exposure groups. **(D, E)** Anxiety-like behavior as assessed by elevated plus maze of 3-6-month-old male and female hAPOE3/4-KI mice (*n* = 8 per group) exposed to PS-NMPs (red), as compared to controls (black). No significant differences in **(D)** percent bouts in open arms and **(E)** percent distance in open arms were found between genotype or exposure groups. Three-way ANOVA to determine the interaction between APOE genotype, sex, and PS-NMPs exposure: **(A)** genotype x sex x exposure p=0.9887, **(B)** genotype x sex x exposure p=0.8981, **(C)** genotype x sex x exposure p=0.8108, **(D)** genotype x sex x exposure p=0.2243, **(E)** genotype x sex x exposure p=0.6336. Sex-specific significances were determined by two-way ANOVA with post-hoc analyses: **(A-C)** no significances**, (D_Male_)** genotype p=0.082, **(D_Female_)** not significant **(E_Male_)** not significant, **(E_Female_)** interaction p=0.041.

In order to determine if exposure to PS-NMPs affected memory in APOE3 and APOE4 h-KI mice, we used the Y-maze and novel object recognition (NOR) assay (Fig. 3). To assess short-term spatial memory using Y-maze, animals explored two arms of the maze while the third was closed off, were then returned to their home cages, followed by a second round of exploration with access to the previously closed off, “novel” arm. We found no significant differences in distance or bouts in the novel arm between APOE genotypes or PS-NMPs exposure groups of either sex (Fig. 3A, B). NOR was then used to assess recognition memory in APOE3 and APOE4 h-KI control and PS-NMPs exposed mice. Animals explored two objects for 30 minutes, returned to their home cage for 2 hours while one object was replaced with a novel object, and were then introduced to both objects for another 30 minutes while movements were monitored in x-, y-, and z-directions. Discrimination index (DI) was calculated as time spent between the novel and familiar objects during the exploratory phase of the test (3 minutes). DI was also calculated for latency to the novel object and for number of bouts with the novel object. No significant differences in measures of recognition memory following PS-NMPs exposure regardless of APOE genotype were found in male mice. We also did not observe any significant impairments in recognition memory following PS-NMPs exposure in female APOE3 mice. We did, however, find marked different in recognition memory in female APOE4 mice exposed to PS-NMPs. These mice spent significantly less time with the novel object (Fig. 3C), were much slower to approach the novel object (Fig. 3D), and also showed fewer, although not significant, approaches to the novel object (Fig. 3E).

**Figure 3.**
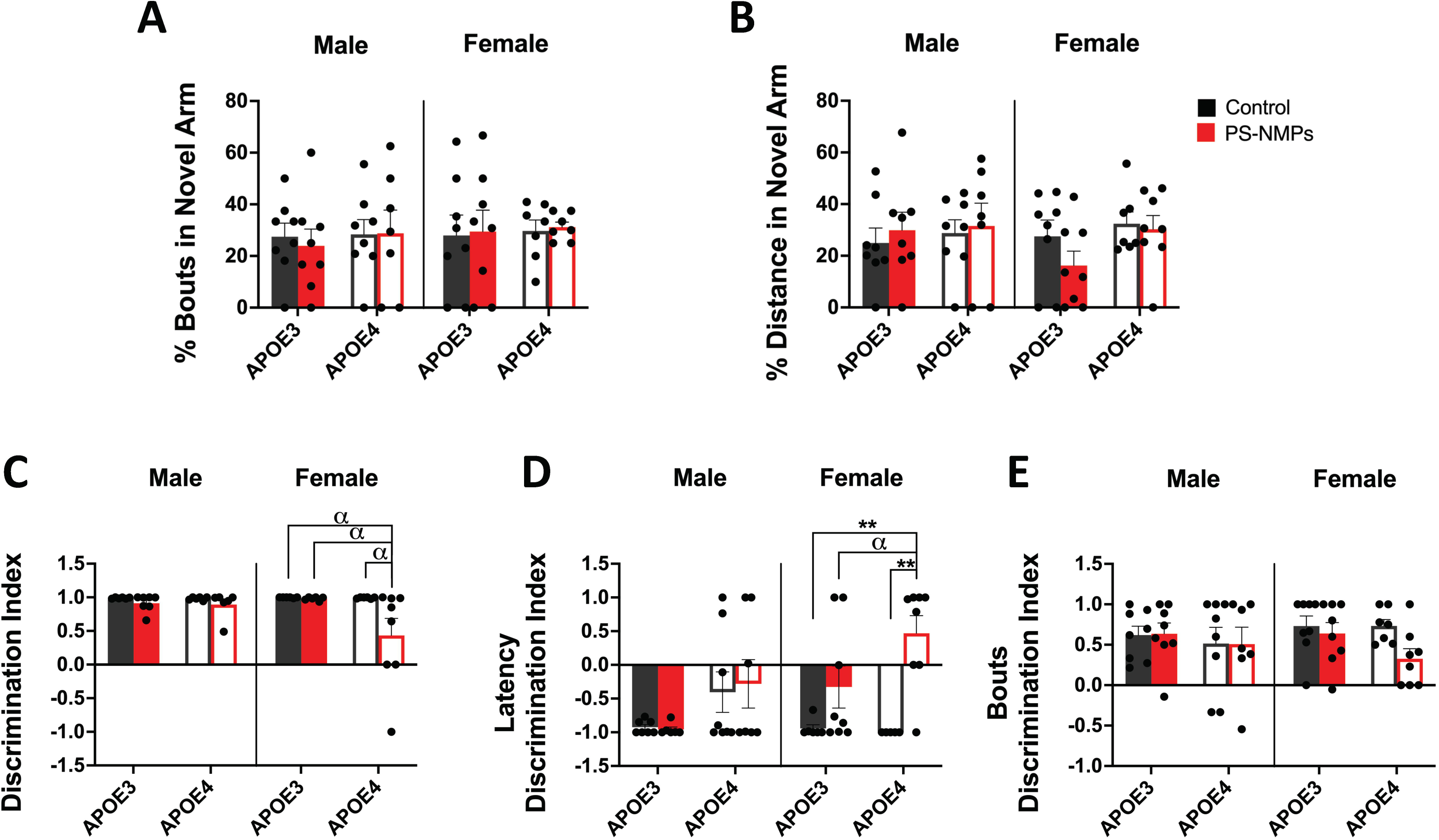
Effect of APOE genotype and exposure to PS-NMPs on memory. **(A, B)** Short-term memory of 3-6-month-old male and female hAPOE3/4-KI mice (*n* = 8 per group) exposed to PS-NMPs (red), as compared to controls (black). No significant differences in **(A)** percent bouts in novel arm and **(B)** percent distance in novel arm of Y-maze were found between genotype or exposure groups. **(C-E)** Recognition memory of 3-6-month-old male and female hAPOE3/4-KI mice (*n* = 8 per group) exposed to PS-NMPs (red), as compared to controls (black). Female APOE4 mice exposed to PS-NMPs exhibited a significant decrease in **(A)** discrimination index and a significant increase in **(B)** discrimination index of latency in novel object recognition task, as compared to APOE3 counterparts as well as APOE4 controls. Three-way ANOVA was used to determine the interaction between APOE genotype, sex, and PS-NMPs exposure: **(A)** genotype x sex x exposure p=0.8302, **(B)** genotype x sex x exposure p=0.5180, **(C)** genotype x sex x exposure p=0.1260, **(D)** genotype x sex x exposure p=0.3317, **(E)** genotype x sex x exposure p=0.4842. Sex-specific significances were determined by two-way ANOVA: **(A)** no significances**, (B_Male_)** not significant, **(B_Female_)** genotype p=0.099, **(C_Male_)** PS-NMPs exposure p=0.097, **(C_Female_)** genotype p=0.098, PS-NMPs exposure p=0.084, **(D_Male_)** genotype p=0.026, **(D_Female_)** PS-NMPs exposure p=0.0004, **(E_Male_)** not significant, **(E_Female_)** PS-NMPs exposure p=0.050, with post-hoc analyses denoted by ** *p* < 0.01 and α *p* < 0.10 as trending.

### Alterations in Inflammatory Markers due to PS-NMPs Exposure

Using fluorescence microscopy, we were able to detect the presence of the 2 µm PS-NMPs in brains of both APOE3 and APOE4 exposed mice as previously done (Gaspar et al., 2023) (Fig. 4). The 0.1 µm particles are below the detection limit in tissue sections using this technique; however, in *in vitro* experiments, we observed similar accumulation of both 2 µm and 0.1 µm particles within the cells, localized around the nuclear envelope following exposure (Gaspar et al., 2023).

**Figure 4.**
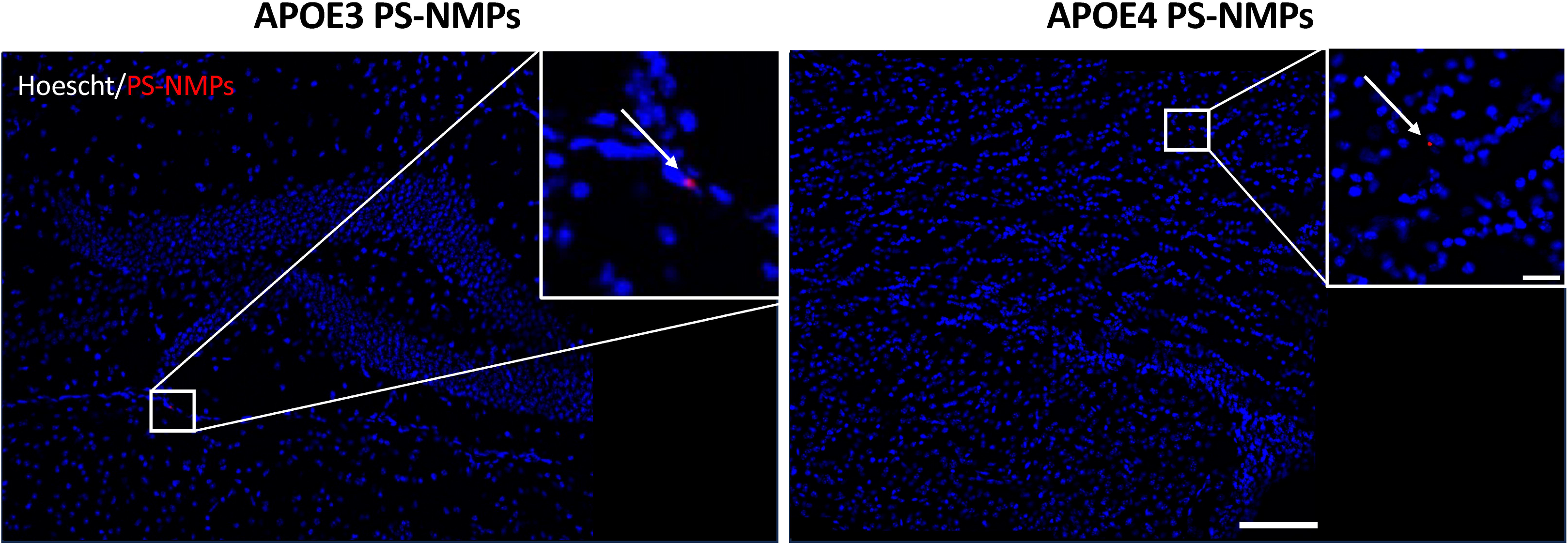
Accumulation of PS-NMPs in brains from APOE3 and APOE4 mice. Representative images from APOE3 and APOE4 h-KI mice (*n* = 4 per group) showing presence of red fluorescent PS-NMPs (red) in hoescht (blue)-stained brain tissue after acute (3 weeks) exposure to PS-NMPs. Scale bar = 100 μm, 20 μm.

To correlate the marked changes in cognition and memory in response to NMPs exposure with tissue histopathology, we evaluated glial fibrillary acidic protein (GFAP), a marker of activated astrocytes using both western blot and immunohistochemistry (Fig. 5). Using these techniques, we observed significant decreases in GFAP expression in APOE3 and APOE4 females following PS-NMPs exposure, with the effects moderately more pronounced in APOE4 mice (Fig. 5D-F). These findings are consistent with our previous results in female C57BL/6J mice similarly exposed to PS-NMPs (Gaspar et al., 2023). Interestingly, in males we observed that GFAP expression decreased in APOE3 but not in APOE4 mice following PS-NMPs exposure. We did, however, note that male APOE4 control mice exhibited decreased GFAP expression compared to APOE3 controls, a trend that was not observed in female mice (Fig. 5A-C). We also identified a seemingly decrease in ionized calcium-binding adaptor molecule 1 (IBA1) expression, a marker for microglia, following PS-NMPs exposure independent of sex and APOE genotype using fluorescent immunohistochemistry (Fig. 5C, F).

**Figure 5.**
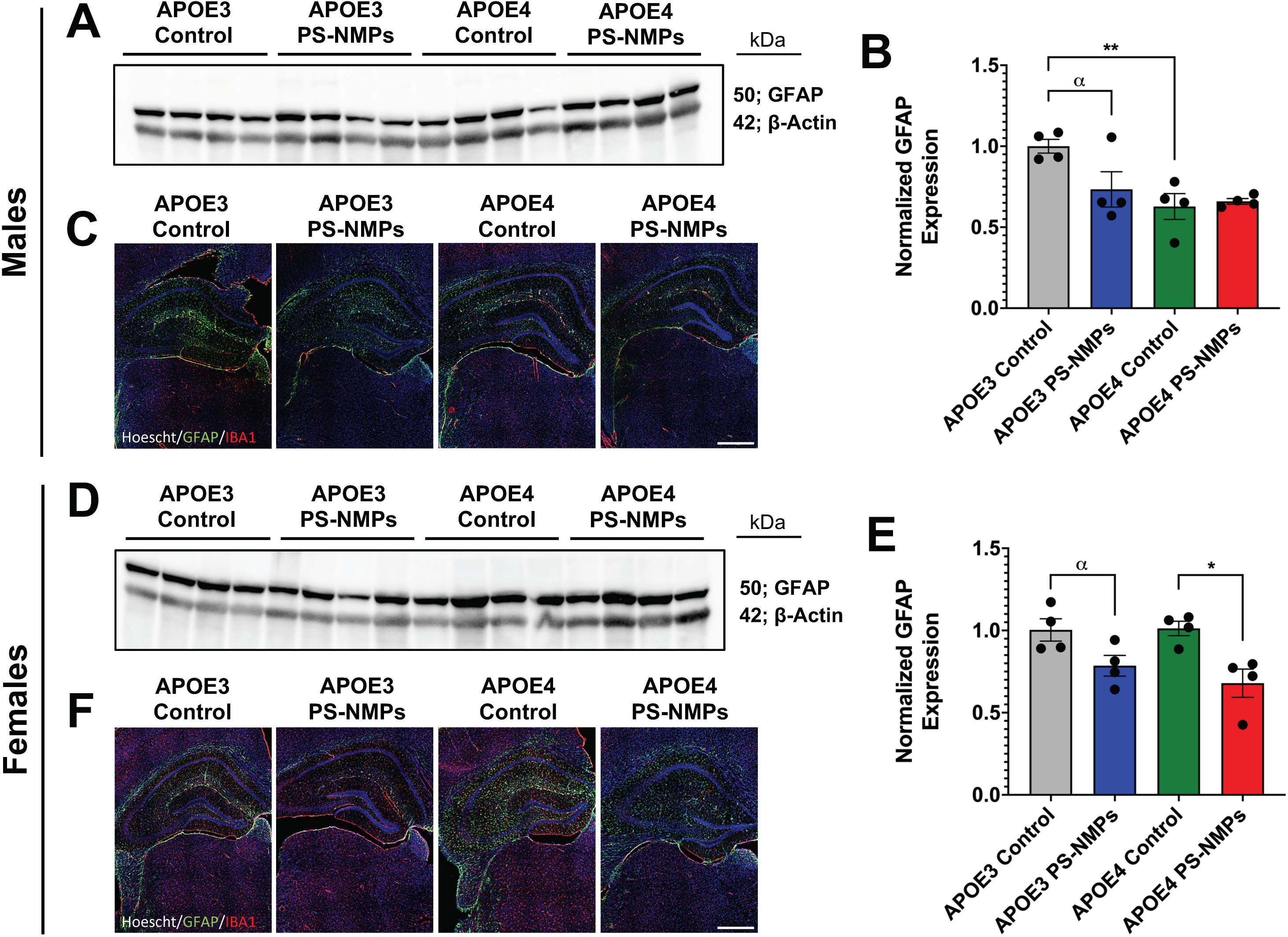
PS-NMPs altered GFAP expression in brains from male and female APOE3 and APOE4 h-KI mice. **(A, B)** Western blot analysis (*n* = 4 per group) revealed decreased GFAP protein levels in male APOE3 mice following PS-NMPs exposure as well as in male APOE4 mice compared to APOE3 controls. No significant differences in GFAP levels were detected between exposure groups in male APOE4 mice. **(C)** Representative images of brain hippocampal sections with hoescht (blue)-staining showing alterations in GFAP (green) and IBA1 (red) following PS-NMPs exposure and between APOE genotypes in male mice (*n* = 4 per group). **(D, E)** Western blot analysis (*n* = 4 per group) revealed decreased GFAP protein levels in both female APOE3 and APOE4 mice following exposure to PS-NMPs. **(F)** Representative images of brain hippocampal sections with hoescht (blue)-staining showing alterations in GFAP (green) and IBA1 (red) following PS-NMPs exposure and between APOE genotypes in female mice (*n* = 4 per group). Significances were determined by unpaired t-tests **(B, E)**, denoted by * *p* < 0.05, ** *p* < 0.01, and α *p* < 0.10 as trending. Scale bars = 100 µm.

### Evaluation of CYP1A1 Expression

To better understand how PS-NMPs might be processed in the body, we performed fluorescent immunohistochemistry and Western blot analysis to determine the protein expression of CYP1A1, an enzyme known largely for its role in metabolizing environmental polycyclic aromatic hydrocarbons (Shimada and Fujii-Kuriyama, 2004)(Chen et al., 2021). In female mice, we found an increasing trend in CYP1A1 protein levels with APOE4 status and exposure to PS-NMPs (Fig. 6E-G). In male mice, however, we observed a decrease in CYP1A1 protein in APOE3 mice exposed to PS-NMPs and no significant changes in APOE4 mice following exposure (Fig. 6A-C). To further investigate these results, we performed qPCR to measure mRNA expression of CYP1A1. In female mice, we observed mild increases in CYP1A1 mRNA expression following exposure to PS-NMPs (Fig. 6H); however, we observed a decreasing trend in CYP1A1 mRNA expression with APOE4 status and exposure to PS-MPs in male mice (Fig. 6D).

**Figure 6.**
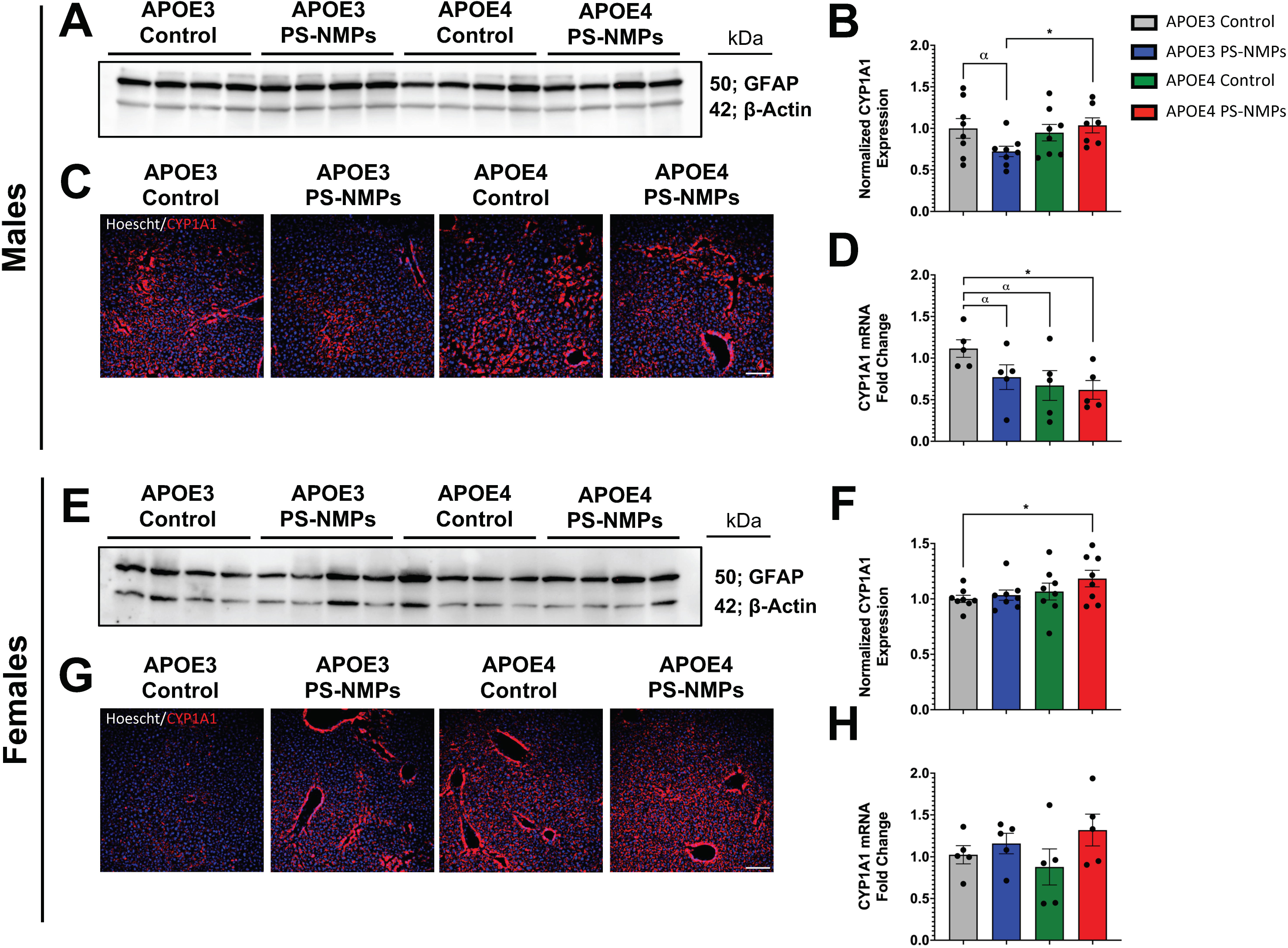
PS-NMPs altered CYP1A1 protein and mRNA expression in livers from male and female APOE3 and APOE4 h-KI mice. **(A, B)** Western blot analysis (*n* = 4 per group) revealed decreased CYP1A1 protein levels in male APOE3 mice following PS-NMPs exposure. No significant differences in CYP1A1 were detected between exposure groups in male APOE4 mice. **(C)** Representative images of liver sections counterstained with hoescht (blue) showing alterations in CYP1A1 protein levels following PS-NMPs exposure and between APOE genotypes in male mice (*n* = 4 per group). **(D)** Measurement of CYP1A1 expression via Taqman qPCR showed significant decreases in mRNA levels with PS-NMPs exposure and APOE4 status in male mice (*n* = 5 per group). **(E, F)** Western blot analysis (*n* = 4 per group) revealed increased CYP1A1 protein levels in female mice following exposure to PS-NMPs and with APOE4 status. **(G)** Representative images of liver sections counterstained with hoescht (blue) showing alterations in CYP1A1 protein levels following PS-MPs exposure and between APOE genotypes in female mice (*n* = 4 per group). **(H)** Measurement of CYP1A1 expression via Taqman qPCR showed moderate increases in CYP1A1 mRNA levels following PS-NMPs exposure in both APOE3 and APOE4 female mice (*n* = 5 per group). Significances were determined by unpaired t-tests **(B, D, F, H)**, denoted by * *p* < 0.05, and α *p* < 0.10 as trending. Scale bars = 100 µm.

## Discussion

As global plastic production and pollution continue to rapidly rise, nano- and micro-plastics pose an increasing threat to environmental and human health. Previous studies have already demonstrated that exposure to NMPs can induce inflammation (Pulvirenti et al., 2022)(Luo et al., 2022)(Gaspar et al., 2023), alter cognition (Gaspar et al., 2023)(Santos et al., 2022), diminish reproductive function (Inam, 2025)(Urli et al., 2023), and result in NMPs bioaccumulation (Gaspar et al., 2023)(Li et al., 2024), amongst other adverse outcomes. In our previous study (Gaspar et al., 2023), we found that acute (3 weeks) exposure to 0.1 and 2 µm PS-NMPs significantly altered behavioral performance and modulated expression of immune markers in both liver and brain tissue in young and old female C57BL/6J mice. We additionally detected the presence of PS-NMPs in every tissue examined, including brain. Based on our findings from this study, we sought to explore how such effects from NMPs exposure might interplay with genetic risk factors and thus contribute to disease onset.

Environmental toxins, such as heavy metals and pesticides, have previously been explored for their contributions to the progression of Alzheimer’s disease (Bakulski et al., 2020)(Yan et al., 2016); however, little work has been done to evaluate how NMPs may impact the development of AD and AD-associated symptoms. To begin addressing this question, we thought to investigate if PS-NMPs could alter cognition of APOE3 and APOE4 humanized knock-in mice, since APOE4 is one of the strongest known genetic risk factors for the development of AD, although there is no clear mechanism for how this may occur. Theories include APOE4 altering lipid transport and metabolism in the brain (“Study reveals how APOE4 gene may increase risk for dementia,” 2021)(Yin, 2023), reducing clearance of amyloid plaques (Prasad and Rao, 2018)(Liu et al., 2017), and contributing to increased levels of neuroinflammation (Parhizkar and Holtzman, 2022)(Dias et al., 2025). Despite all of these, however, it is clear that the presence of APOE4 alone is not sufficient to cause AD and there must be other contributing factors.

In the present study, we exposed 3-6 months old female and male APOE3 and APOE4 h-KI mice to a 1:1 volume mixture of 0.1 and 2 µm pristine spherical fluorescent PS-NMPs for 3 weeks via drinking water (Fig. 1A) to assess if acute NMPs exposure in mice carrying the APOE4 allele may promote the development of AD symptoms. After 3 weeks of NMPs exposure, we first conducted a battery of behavioral assays including open-field, light/dark preference, elevated plus maze, Y-maze, and novel object recognition. Results from light/dark preference, EPM, and Y-maze showed no significant changes in behavior following exposure to PS-NMPs in female and male APOE3 and APOE4 mice (Fig. 2 and 3A, B). In open-field, however, we observed a striking increase in the percentage of time that male APOE4 mice exposed to PS-NMPs spent and rested in the center of the arena (Fig. 1B-D), suggesting cognitive dysfunction. Conversely, in the NOR assay we observed marked memory impairments in female APOE4 mice exposed to PS-NMPs as measured by the discrimination index (Fig. 3C, D). It is notable that these effects were not observed in APOE3 mice of either sex, suggesting that the combination of APOE4 and PS-NMPs exposure resulted in these behavioral alterations. Additionally, the behavioral changes observed in the APOE4 mice following PS-NMPs exposure are sex-dependent. While the exact pathway for this remains unknown, possibilities for this discrepancy include differing brain compositions in males and females, hormonal differences, and differing lipid profiles in the brain (Lopez-Lee et al., 2024)(Gur et al., 1999)(Guo et al., 2022). These factors are all likely contributing to the apathy-like behavior only exhibited by the male APOE4 mice exposed to PS-NMPs in the open-field assay, as well as the decreased recognition memory predominantly exhibited by the female APOE4 mice following exposure. Interestingly, these sex-dependent differences strongly relate to human clinical symptoms of AD, with men more often presenting with symptoms such as apathy and impaired motor control and women often presenting with symptoms related to memory loss (Liampas et al., 2022)(Dolphin et al., 2023).

Following these behavioral observations, we then set out to explore how PS-NMPs may be altering immune markers in the brain. In our previous study (Gaspar et al., 2023), we observed a marked decrease in GFAP expression following PS-NMPs exposure, particularly in younger animals. We postulated that such a decrease may be indicative of astrocyte atrophy that may occur early in AD prior to the hypertrophy that is characteristic of increased neuroinflammation. In both APOE3 and APOE4 female mice, we once again observed decreased GFAP expression following PS-NMPs exposure (Fig. 5D-F). Interestingly, this trend was also observed in male APOE3, but not APOE4, mice exposed to PS-NMPs (Fig. 5A-C). We did, however, find that male APOE4 control mice already expressed decreased levels of GFAP, thus PS-NMPs exposure combined with the APOE4 allele may be driving astrocyte hypertrophy rather than atrophy in APOE4 males. Additionally, we found that there was a seemingly general trend towards decreased IBA1 expression in all groups following PS-NMPs exposure (Fig. 5C,F) indicating a decrease in microglia. Interestingly, a recent report from another group using PS-MPs of a similar size (2.5 µm) showed that exposure induced microglial pyroptosis in mice, which may explain the decreased IBA1 levels we observed in the present study (Wang et al., 2024).

NMPs have also been shown to interact with every system of the body and may potentially have peripheral effects that can be impacting cognition and overall brain health. For example, we previously observed increased expression of inflammatory cytokine TNF-α and environmental alarmins S100a8 and S100a9 in liver tissue following PS-NMPs exposure (Gaspar et al., 2023). We specifically chose to examine the effects in liver for its major role in blood detoxification, given that NMPS have been shown to travel via the bloodstream. While we have confirmed increased inflammation of the liver following NMPs exposure, we were interested in exploring another major liver component, cytochrome P450s (CYPs). CYPs represent the major drug metabolizing enzymes in the body and handle a vast majority of the metabolism and detoxification of many xenobiotics (Esteves et al., 2021)(Guengerich, 2021). In particular, we chose to focus on CYP1A1 for its role in metabolizing polycyclic aromatic hydrocarbons and its previously reported robust response to other environmental toxins, such as 2,3,7,8- Tetrachlorodibenzo-p-dioxin (TCDD) (Chen et al., 2021)(Patrizi and Siciliani de Cumis, 2018). To our knowledge, this is the first study to explore the response of a CYP enzyme to NMPs, which may provide valuable insights into NMPs toxicity and processing within the body. In female mice, we observed that CYP1A1 expression exhibited a trending increase with PS-NMPs exposure and APOE4 presence (Fig. 6E-H). Interestingly, we found that CYP1A1 expression decreased in male APOE3 mice and was not significantly altered in male APOE4 mice, following PS-NMPs exposure (Fig. 6A-D). While the mechanisms underlying these observations are undoubtedly complex, a simplistic proposed theory is that PS-NMPs may induce increased expression of CYP1A1, which in turn increases oxidative stress. Other studies have reported that excessive oxidative stress should negatively feedback to inhibit enzymatic activity and/or transcription of CYP1A1 (Barouki and Morel, 2001)(Nebert et al., 2000). This could explain the decrease in CYP1A1 expression in male APOE3 mice following PS-NMPs exposure. It has also been theorized that estrogen can have an antioxidant effect (Xiang et al., 2021)(Bellanti et al., 2013), which may thus minimize oxidative stress in female mice allowing for increased CYP1A1 levels following PS-NMPs exposure. As for male APOE4 mice, it remains unclear why CYP1A1 expression was not modulated following PS-NMPs exposure. One possible theory, that would involve additional experimentation to support, is that the presence of APOE4 is somehow modulating CYP metabolism by inhibiting the negative feedback loop that is activated in an excess of oxidative stress to reduce CYP1A1.

Overall, the results from this study demonstrate that exposure to PS-NMPs can alter cognition and memory as well as immune markers in the brain predominantly in mice carrying the APOE4 allele, the strongest genetic risk factor for developing AD. Our results also suggest that PS-NMPs may be involved in the induction of CYP1A1, which could potentially be contributing to their toxicity. These findings further highlight the complicated nature of AD development, which can be influenced by numerous factors, including sex, APOE genotype, exposure to environmental toxins, as well as other factors not explored here. It has previously been shown that humans with diseases such as AD and liver cirrhosis have on average higher MPs accumulation (Horvatits et al., 2022)(Nihart et al., 2025), but what is unclear is whether MPs accumulation increases as a result of the disease or if the disease is in part triggered by increased MPs accumulation. Such questions are nearly impossible to answer in human tissues; however, studies such as this are exceedingly important and can begin to provide a insight into potential pathways and interactions that may result in disease symptomatology following NMPs exposure. There is still significant work that must be done to better understand the interconnectedness between NMPs exposure, APOE genotype, sex, and CYP metabolism and how all of these factors may be contributing to AD-like symptoms. Future studies should further explore parameters such as accumulation of NMPs in APOE3 versus APOE4 individuals, how differences between sexes including hormone levels and brain matter composition may alter NMPs exposure outcomes, and the response of a wider range of CYPs to NMPs. Such studies will be pivotal in shaping our understanding of NMPs toxicity and how they may be a critical driver in the development of neurodegenerative diseases.

## Author Contributions

L.G., G.C., and J.M.R. designed the experiments. L.G., S.B, and H.T.W. performed the experiments, and L.G. analyzed the data. L.G., G.C. and J.M.R. wrote the manuscript and all authors have read and agreed to the published version of the manuscript.

## Acknowledgements

This research was supported by the Rhode Island Medical Research Foundation (JMR), the Plastics Initiative (JMR) at the University of Rhode Island, the College of Pharmacy (GC, JMR) at the University of Rhode Island, the George & Anne Ryan Institute for Neuroscience (GC, JMR) at the University of Rhode Island, the University of Rhode Island Faculty Development Grants (JMR), and the Rhode Island Institutional Development Award (IDeA) Network of Biomedical Research Excellence from the National Institute of General Medical Sciences of the National Institutes of Health under grant number P20GM103430 (JMR).

## Institutional Review Board Statement

The animal study protocol (#AN1920-020) was approved by the Institutional Review Board of University of Rhode Island, under J.M.R. and G.C.

## Data Availability Statement

Data generated from this study are available upon request.

